# Cleanet: robust doublet detection in cytometry data based on protein expression patterns

**DOI:** 10.1101/2025.01.09.632259

**Authors:** Matei Ionita, Michelle L. McKeague, Mark M. Painter, Divij Mathew, Ajinkya Pattekar, Damian Maseda, E. John Wherry, Allison R. Greenplate

## Abstract

Flow and mass cytometry experiments are essential for profiling immune cells at single cell resolution. Better understanding of human immunology increasingly involves analyzing studies at the scale of hundreds or thousands of samples, with data analysis a significant bottleneck. This trend increases the demand for automated analysis methods. In particular, a common preprocessing step in cytometry data analysis is distinguishing single cells from doublets (or multiplets), events in which two (or more) cells pass simultaneously through the detector. Typically, doublets are identified on two-dimensional density plots, using their high measured values for DNA intercalators (mass cytometry) or scattering channels (flow cytometry). Despite its popularity, this bivariate gating method is sometimes imprecise: for example, we show that bivariate gating of mass cytometry data can mistake single eosinophils for doublets, due to their high DNA content. Taking inspiration from methods already used in single cell transcriptomics, but not in the cytometry community, we propose an alternative approach. Our method, called Cleanet, first simulates doublet events, then identifies true events with protein expression similar to the simulated doublets. This simple method is completely automated and detects both homotypic and heterotypic doublets. We validate it in datasets acquired with mass and flow cytometry; moreover, we verify with imaging flow cytometry that events predicted to be doublets truly consist of multiple cells. Cleanet can also classify doublets based on their component cell types, which potentially enables the study of cell-cell interactions, mining extra information out of doublet events that would otherwise be discarded. As a proof of concept, we demonstrate that Cleanet can detect a treatment-specific increase in interactions between two cell lines. By automating doublet detection and classification, we aim to streamline the data analysis in large cytometry studies and provide a more accurate picture of both immune cell populations and cell-cell interactions.

## Introduction

Flow and mass cytometry studies are becoming increasingly complex as they involve higher dimensional data and larger sample sizes. In order to scale to thousands of samples, studies would benefit from automating as much of the analysis tasks as possible. In particular, data cleaning is a good target for automation, both because it is important to ensure minimal variability in a task that influences all downstream analysis, and because it requires less biological interpretation than other downstream tasks. To this end, multiple automated methods have been proposed to remove anomalous events: FlowFP^1^, flowAI^2^, flowClean^3^, PeacoQC^4^, flowCut^5^.

This work focuses on the specific problem of detecting multiplets, two or more cells which pass through the detector at the same time and are registered as a single event. In cytometry data, doublet detection is important in order to avoid spurious phenotypes or biased cell type proportions. Traditionally, doublets are removed by manual gating on bivariate density plots: events with high DNA intercalator in mass cytometry, and events with high scattering area for a given scattering height in the case of flow cytometry. The R package PeacoQC has automated the bivariate scattering gate for flow cytometry, by flagging events whose ratio of scattering area to scattering height is more than a number of median absolute deviations above the median.^4^ More recently, SingletSeeker proposed distinguishing debris, singlets and multiples using density-based clustering on scattering channels, or on imaging parameters if available.^6^

In addition, other computational methods have been proposed to automate bivariate gating more generally. FlowDensity finds peaks and valleys in univariate or bivariate densities in order to determine gate locations.^7^ UNITO uses image registration and a convolutional neural network model to learn the position of gates from training examples and predict it in new samples.^8^ Elastigate also uses image registration, but then transforms testing images to match training ones and transfers the gates.^9^ These or other methods for automatic gating can be applied to doublet detection. However, they are limited by the fact that singlets and doublets are not always peaks separated by a valley, or by the availability of training data.

Moreover, a limitation of all existing methods is that they make inferences about doublets based on either a bivariate plot or a small number of imaging parameters, ignoring information contained in the protein expression channels. In many cases, the singlet and doublet distribution overlap somewhat along the channels traditionally used for singlet gates, even though they may be well separated in multivariate space. For example, in flow cytometry, the “width” and “area” scattering channels have higher values for doublets, but most of the time there is a continuum of values rather than a clear separation. This problem is exacerbated by cell type heterogeneity, as cell types of different size and complexity have different expected values for scattering or DNA intercalator channels. Intuitively, a multivariate method can improve the classification of edge cases by looking for events which express marker proteins from multiple lineages (heterotypic doublets) or express approximately twice as much protein as other phenotypically similar cells (homotypic doublets).

To formalize this intuition about the benefits of multivariate analysis, the experience of the single cell transcriptomics community is a useful guide. scRNA-seq data has many more channels than cytometry data, but each channel is individually noisier^10^. Accordingly, that community has developed methods for doublet detection that use the full multivariate gene expression data, such as DoubletFinder^11^, Scrublet^12^, DoubletDecon^13^, scDblFinder^14^ and Chord^15^. In this manuscript, we take inspiration from DoubletFinder and Scrublet, which proposed a simple and elegant idea: generate synthetic doublets by combining the expression of pairs of single cells, then look for events in the original data which are similar to the synthetic doublets.

Cleanet is an implementation of this approach adapted to cytometry data, with modifications to account for technical differences between the experimental platforms. To the best of our knowledge, this is the first time that a method for doublet detection based on multivariate expression data has been proposed for flow or mass cytometry. We validate Cleanet on both mass and flow cytometry datasets, showing that it performs well even in the presence of technical artifacts. Moreover, in rare cases where Cleanet predictions differ significantly from those of a manual approach, we argue that Cleanet better reflects cell type protein expression patterns. Looking for an objective ground truth, we also use ImageStream brightfield images of cellular events and check that they validate Cleanet predictions of singlets and doublets.

Aside from automation, another benefit of Cleanet is the native ability to extend a given classification of singlets into a classification of the doublets. For example, if the singlets are classified as lymphocytes or monocytes using any manual or automated method, Cleanet extends this into a classification of doublets as lymphocyte-lymphocyte, lymphocyte-monocyte or monocyte-monocyte. This is of interest when some doublets arise from real cell-cell interactions, not just from technical factors, and the proportions of doublet types can serve as an immune feature or biomarker alongside the proportions of singlets^16^. As a proof of concept, we use a cell line experiment in which treatment with Rituximab is expected to increase the rate of interactions between the Raji and THP1 cell lines. Cleanet detects a treatment-dependent increase in Raji-THP1 heterotypic doublets.

## Results

### Method overview and comparison to single cell transcriptomics

Cleanet uses single cell events from each data file to simulate the expected distribution of protein expression in doublets from that file. Because of the curse of dimensionality, it is difficult to model the multivariate density function of doublets in high dimensional data. However, it is easy to sample from this distribution. Indeed, sampling pairs of cells at random and adding the protein expression values from the two cells in each pair is sufficient. (Figure 1A) Cleanet then looks for events in the data file for which at least a third of nearest neighbors are simulated doublets, and predicts that they are the true doublets. (Figure 1B)

**Figure 1.**
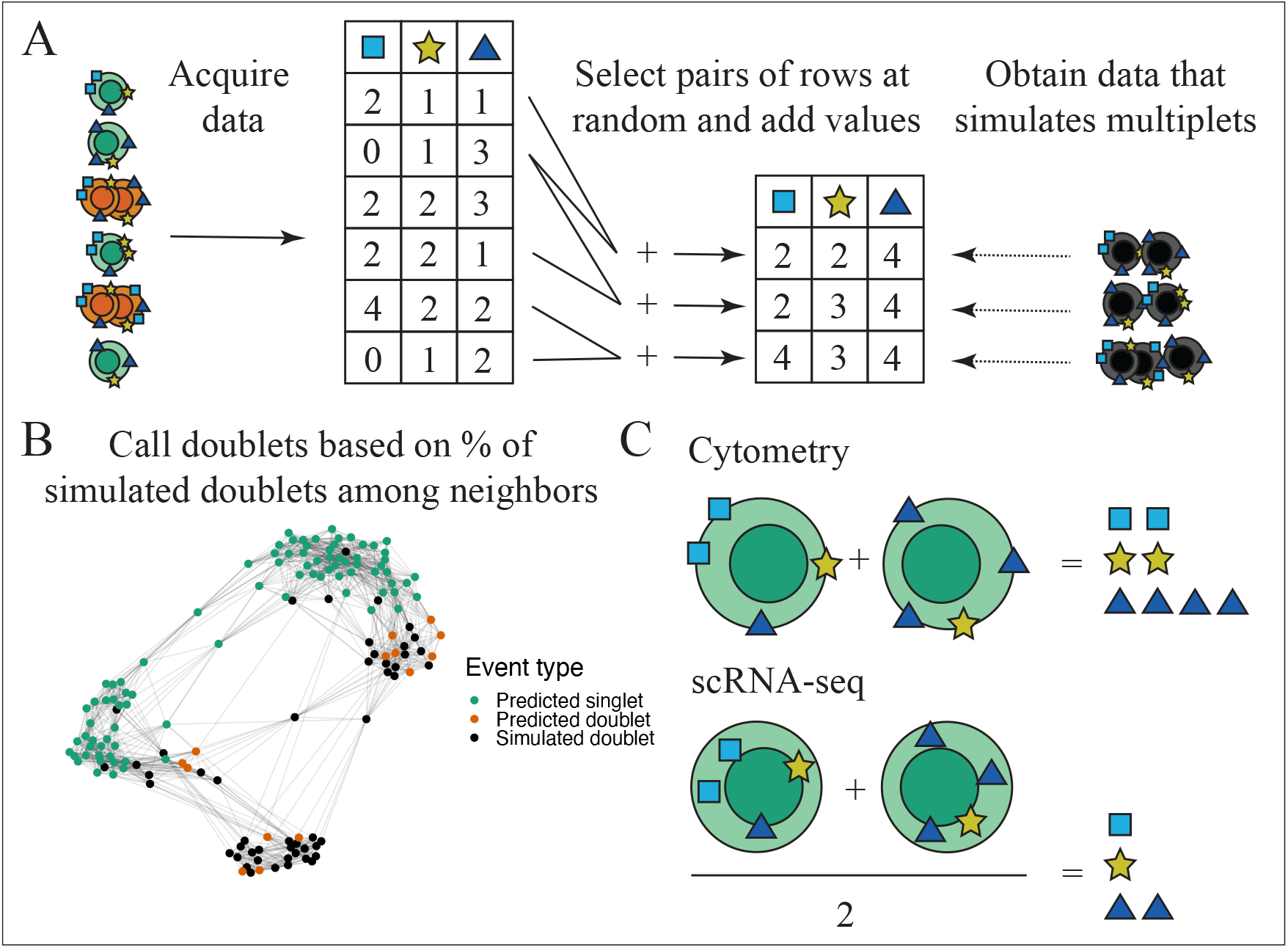
(A) Cleanet begins by augmenting data with simulated doublet events. These are obtained by sampling pairs of true events and summing the protein expression of the two components in each pair. (B) Nearest neighbor graph constructed by Cleanet for augmented dataset (real and simulated events together). Those events (red) which have a high share of simulated doublets (black) among their 15 nearest neighbors are classified as doublets. All other events (green) are classified as singlets. (C) Mechanism for doublet simulation in Cytometry versus scRNA-seq. A cytometry doublet event records the sum of signals from individual cells, whereas a scRNA-seq doublet event records the average gene expression, because of limited sequencing depth.

The current work adapts this idea from the setting of single cell transcriptomics to that of cytometry. This adaptation requires some technical modifications, the most consequential of which is the way we model a doublet from its constituent parts. In current scRNA-seq protocols, the sequencing depth is limited to a number much smaller than the total amount of transcripts present in the cell.^17^ Because of this, the gene expression of a doublet event is best modeled as the average expression of the single cells. In contrast, cytometry assays aim to saturate the target proteins with antibody probes, so the protein expression of a doublet is best modeled as the sum of protein expression in single cells (Figure 1C).

This technical difference between assays has a major impact on the detection of homotypic doublets (those composed of two cells of the same type). In single cell transcriptomics, the average expression of two identical cells looks like a third cell identical to the first two, therefore it is computationally indistinguishable from them. But in cytometry, a homotypic doublet is easily distinguished because it looks like an event with twice as much protein expression as the individual components. Thus, Cleanet performs just as well for homotypic and heterotypic doublets.

### Cleanet detects doublets in mass cytometry data and reduces variability in cell type proportions

We first demonstrate Cleanet on a mass cytometry dataset containing 169 whole blood profiles (Methods). This dataset was chosen because there is significant technical variability between the files, especially in the channels commonly used for single cell gates, making it a challenging test case for a doublet detection algorithm. Moreover, manual annotation of doublets was available for all files, enabling a direct comparison between Cleanet and manual gating.

Predicted doublet status was visualized using bivariate plots of a DNA intercalator (traditionally, doublets are defined as events with high amounts of intercalator, because two cells have more DNA than one) versus CD45 (used to distinguish CD45hi lymphocytes from CD45mid granulocytes). Across files with varying distribution profiles of DNA intercalator staining, Cleanet consistently identified events with high DNA intercalator as doublets and broadly agreed with manual gating (Figure 2A). There were, however, two sources of disagreement. The first was over the precise cutoff between single cells and doublets, especially in files where the DNA intercalator did not completely separate the singlet and doublet distributions. The second was over mid-sized segments of the bivariate plot in some files which were classified as singlets despite having a lot of DNA intercalator.

**Figure 2.**
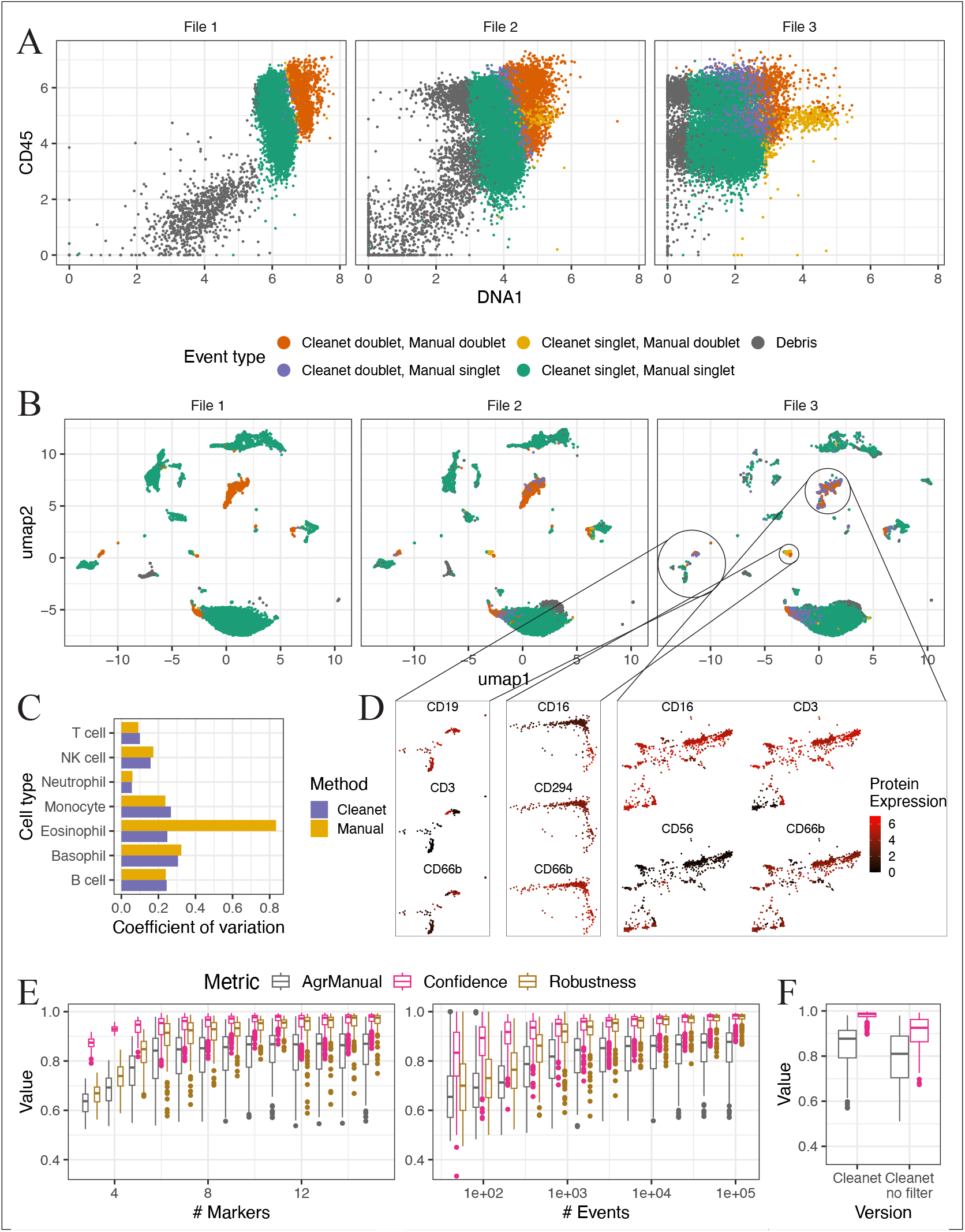
Cleanet distinguishes doublets in CyTOF data. (A) Doublets detected by Cleanet or manual gating in three files, in decreasing order of data quality. In file 3, the DNA intercalator has poor staining, impairing the manual gating of intercalator-high events. (B) Same data shown in UMAP projection of all 32 channels in the panel. (C) Using Cleanet for doublet removal leads to more stable quantification of Eosinophil proportions across 19 technical replicates. Coefficient of variation = standard deviation / mean, lower is better. (D) In files 2 and 3, Cleanet is more consistent than manual gating at labeling CD294hi CD66bhi CD16lo Eosinophils as singlets, and events expressing markers from multiple lineages as doublets. (E) Robustness and agreement with manual gating (objective metric) and Confidence (internal metric) degrade simultaneously and gracefully as the number of events or markers decreases. Boxplots show median and interquartile range over 169 files. (F) Depleting doublets with a filtering function included in the Cleanet package improves both confidence and agreement with manual gating.

To understand these discrepancies, we analyzed the protein expression of the events in question. UMAP dimensional reduction plots overlaid with doublet predictions from the two methods are shown in Figure 2B, and areas of interest overlaid with arcsinh-transformed protein expression values are shown in Figure 2D. Doublets predicted by the automated method are more contiguous in UMAP space, and better cover UMAP islands which express markers from multiple lineages, indicative of heterotypic doublets. This suggests that the cutoffs between singlets and doublets made by Cleanet can be more meaningful than those made by manual gating. Furthermore, the mid-sized segments of Cleanet singlet/Manual doublet events express the eosinophil markers CD66b and CD294, and no other lineage markers. Thus, they are most likely eosinophil singlets, in which high DNA intercalator was measured for reasons that may be biological or technical. Here too, the decision made by Cleanet seems more consistent with the full expression profile of the events than the manual annotation.

We next sought an additional source of validation for the putative eosinophil singlets. Among the 169 files in the dataset, 19 are technical replicates (aliquots from the same blood draw of a healthy donor that were used as day of run controls in the study). As such, we expect cell type proportions in these 19 files to be similar despite technical artifacts, like inconsistent DNA intercalator staining, that make gating challenging. We measured the proportions of seven major cell types, both by a fully manual approach (manual gating following manual doublet removal) and by a fully automated one (unsupervised clustering following Cleanet doublet removal).

Figure 2C shows coefficients of variation (standard deviation divided by mean) for both approaches. Notably, proportions for eosinophils are much less variable using the high-dimensional, automated approach. This is evidence that the bivariate method does not reliably distinguish eosinophils from doublets, at least in this technically challenging dataset. By paying attention to multiple channels at once, Cleanet avoids this pitfall and provides more reliable data cleaning.

Next, we tested Cleanet’s robustness to decreasing the number of events or markers that are available. More specifically, for each of the 169 files, either the number of events or the number of markers was downsampled, and results were compared with both manual gating (AgrManual) and with Cleanet predictions for the entire file (Robustness). In each case, balanced accuracy was computed, to account for the imbalance between abundant singlets and rare doublets. The results show that both metrics degrade gracefully and remain above 80% when as few as 500 events or as few as 6 markers are used (Figure 2E). When downsampling markers, the steepest drop is registered once both DNA intercalator channels are removed, which is consistent with DNA intercalators being the most predictive of doublet status. However, even in the absence of the intercalators, removal of more markers further degrades performance, showing that multiple markers are informative for doublet detection.

Cleanet is also endowed with an internal measure of confidence in its predictions, which can be used in the absence of any external reference to gauge whether Cleanet is a good match for the dataset under consideration. Confidence is defined as the percentage of synthetic doublets which would be correctly classified as doublets by the Cleanet decision criterion. A low confidence value would mean that singlets and doublets are too similar in high dimensional space for quality predictions. This could happen, for instance, when the data has low dimensionality, the markers have poor staining index, or the markers are only expressed on very few cells. In our mass cytometry experiments, confidence starts around 95% and decreases in tandem with robustness as the number of events or markers is decreased (Fig. 2E). To the extent that confidence is higher than agreement with manual gating, this could reflect overconfidence of the model. Alternatively, it could reflect imprecise manual gating: Cleanet made high confidence, low agreement predictions for files with poor DNA intercalator staining, for which manual gating of doublets was difficult (Supplementary Fig. 1B, 1E).

Variable amounts of debris were present in the mass cytometry files, arising from incomplete red blood cell lysis, platelets or other non-immune events. If present in large proportions, debris can hamper Cleanet’s simulation mechanism, because a singlet-debris pair fails to add up to a doublet. The Cleanet package provides functions with visual feedback to deplete debris in the sample before doublet detection (Methods). In the absence of this step, Cleanet performance deteriorates, with the largest decrease in files that contain more than 20% debris (Fig. 2F, Supplementary Fig. 1B, 1C, 1D). Thus, while debris is an important challenge, depleting it to under 10% of total events ensures adequate performance for Cleanet.

### Cleanet detects doublets in flow cytometry data and classifies doublet composition

Cleanet detects doublets based on their high dimensional protein expression, a method that works for flow cytometry just as well as for mass cytometry. To validate Cleanet predictions for fluorescence flow data, we used a 16-color dataset consisting of 126 PBMC samples^18^, publicly available on FlowRepository^19^.

Aside from fluorescence channels, which depend on a complex and possibly noisy interplay of antibodies and fluorochromes, flow cytometry data also provides scattering channels that measure relatively clean optical phenomena. Traditionally, doublets in flow cytometry are defined by manual gating as events having high scattering area for a given value of scattering height. This manual approach has been automated in the PeacoQC package. Since manual gating was not available for the flow data, we compared PeacoQC and Cleanet for doublet detection. As for mass cytometry data, debris was first depleted using a function provided in the Cleanet package before proceeding to doublet identification (Supplementary Fig. 2A).

Doublets identified by the two automated methods agreed to a large extent (Figure 3A). Moreover, when downsampling the 16 available markers in an analogous manner to the mass cytometry experiment, agreement between Cleanet and PeacoQC degraded similarly to Cleanet’s robustness and confidence metrics (Figure 3B). Performance on all metrics was somewhat lower and degraded faster compared to the mass cytometry experiment; we speculate that this is because the flow cytometry dataset contains fewer bimodal lineage markers (like CD3 or CD14) and more markers with dim or infrequent expression (like PD1 or Ki67).

**Figure 3.**
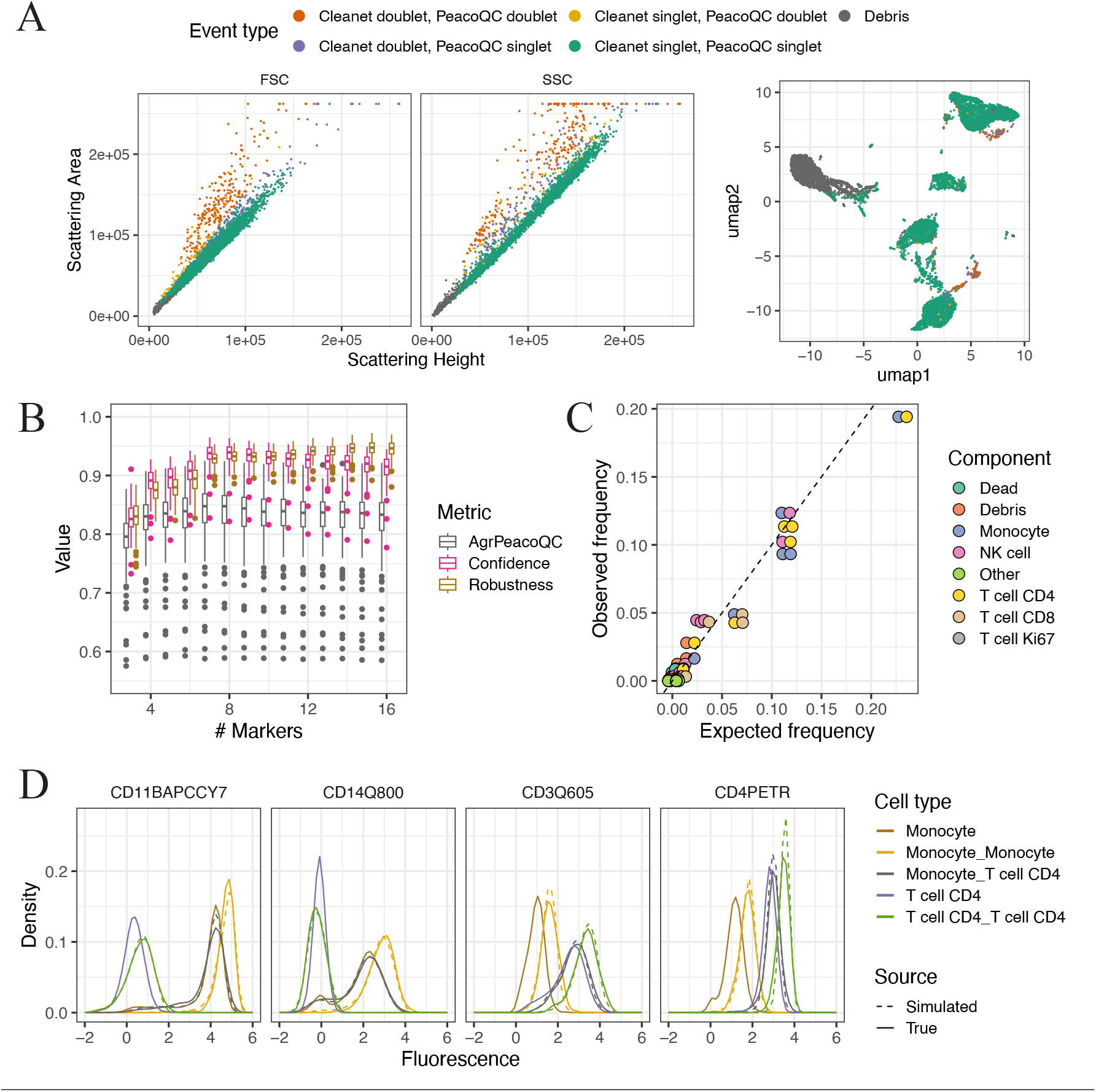
Cleanet distinguishes doublets in flow cytometry data. (A) Cleanet doublet detection agrees with PeacoQC and with the intuition that doublets have high area to height ratio in their front and side scatter profiles. (B) Robustness, Confidence and agreement between Cleanet and PeacoQC decline gracefully as the number of markers is decreased. (C) If provided with cell type labels for the single cells, Cleanet classifies doublets based on their composition. Observed frequency of doublet types in Cleanet output has high correlation the expected frequency based on the abundance of each component. Each pair of points represents a doublet type, and color shows the cell type identity of the two components. (D) Kernel density estimates for the distribution of two monocyte markers (CD11b, CD14) and two T cell markers (CD3, CD4) in Monocytes, CD4 T cells and the three types of doublets they can form. True doublets (solid line) are similarly distributed to simulated doublets (dashed line). Fluorescence is arcsinh-transformed.

As a consequence of using simulated doublets whose composition is known, Cleanet can be extended to also determine the cellular identity of predicted doublets (Methods). A further input necessary for this doublet composition module is a classification of singlets into cell types, which can be provided by the user at any time after the doublet detection stage. For this flow cytometry experiment, FastPhenograph^20,21^ was used to cluster the singlets predicted by Cleanet, then each cluster was manually labeled as one of five cell types (T cell CD4, T cell CD8, T cell Ki67, NK cell, Monocyte), Dead cells or a catch-all category called Other (Supplementary Fig. 2B). Then Cleanet was used to determine the composition of the predicted doublets: for example, they could be T cell CD4 homotypic doublets, or Monocyte-T cell CD4 heterotypic doublets. As a sanity check, we verified that the observed frequency of each predicted doublet type agrees with the expected frequency, based on the product of abundances of the two components (Fig. 3C, Methods). Thus, Cleanet can classify doublets by composition and can flag types of doublets that have unexpected frequency, for biological or technical reasons.

Once cell type labels were available for singlets, true doublets and simulated doublets, the distribution of protein expression in these types of events was compared, focusing on monocytes, CD4+ T cells and the three types of doublets they generate (Fig. 3D). The distributions for true doublets and simulated doublets were very similar, validating the simulation framework used by Cleanet. Moreover, the distributions for homotypic doublets (such as monocyte-monocyte) are shifted towards the right compared to singlets (such as monocyte), providing a visual confirmation that the data is rich enough to distinguish them.

### Validation by imaging cytometry

As a final validation for Cleanet, we turned to Amnis ImageStream data, which captures images of cellular events alongside fluorescent measurements. Cleanet was used to distinguish singlets from doublets based on the fluorescent channels, then the imaging data was used to validate the predictions. The ImageStream dataset measured samples in which the Raji B cell lymphoma and monocytic THP1 cell lines were mixed. One sample was left untreated, while the other was treated with the anti-CD20 monoclonal antibody Rituximab.^22^ Rituximab was expected to increase the rate of interactions between the two cell lines, leading to a higher fraction of heterotypic than homotypic doublets in the treated sample. We sought to test whether this effect could be detected using Cleanet’s capability to classify doublet composition.

Cleanet was used to distinguish doublets using the five available channels (two cell trace channels, two brightfield and one viability channel) and obtained results that were similar to a standard doublet gate using morphology channels (Fig. 4A, 4B). Cleanet reported internal confidence scores of 0.91 and 0.90 for the two samples, which is relatively high given the few available fluorescent channels. Singlets were classified with a bivariate gate based on the fluorescence intensity of the membrane dyes CellTrace Yellow (CTY) and CellTrace Violet (CTV), which labeled Raji and THP1 cells, respectively (Supplementary Fig. 3B). Using this information about singlets, Cleanet was used to classify doublet types as Raji homotypic, THP1 homotypic, or Raji-THP1 heterotypic. The distributions of real and simulated doublets were similar (Fig. 4C).

**Figure 4.**
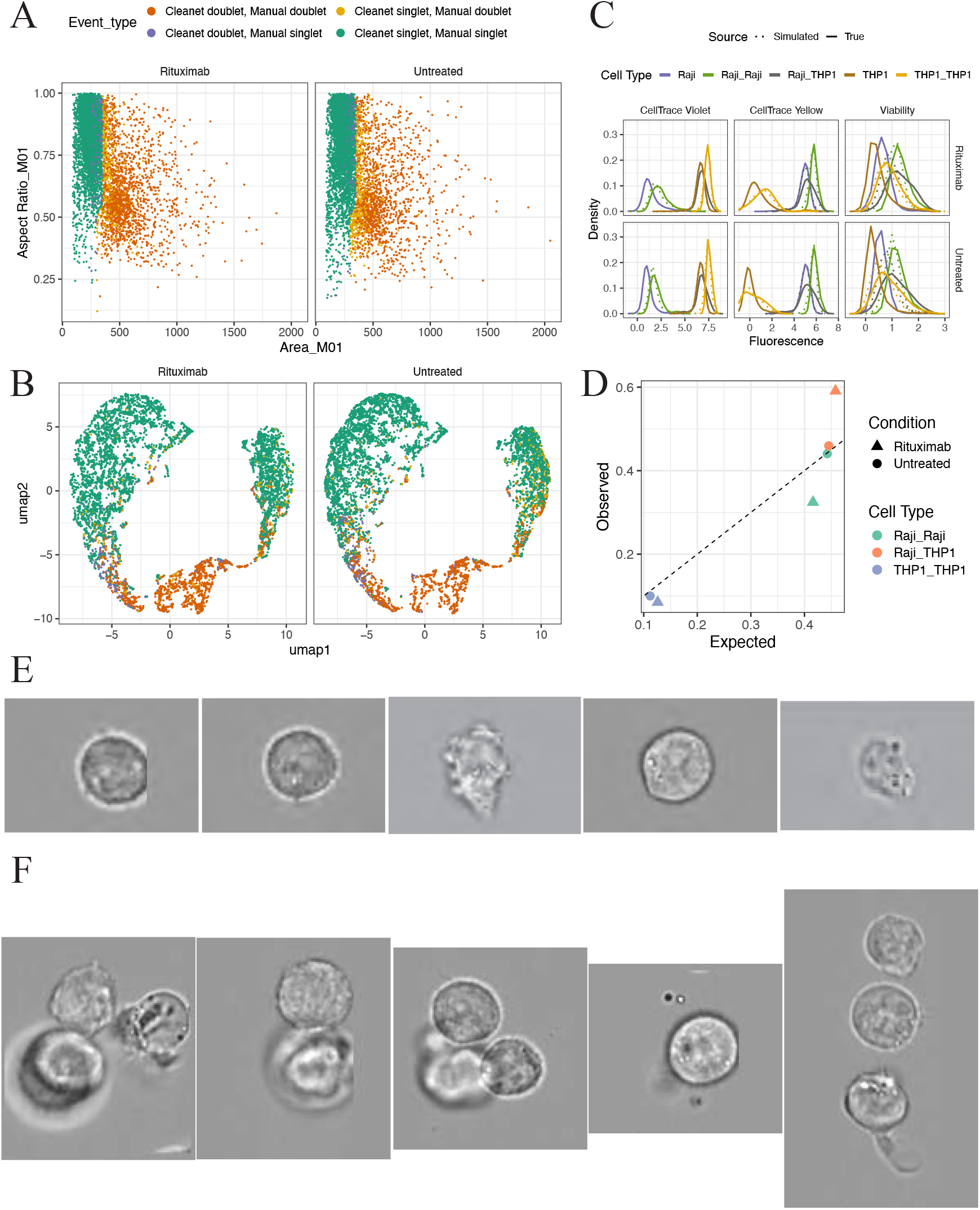
Cleanet distinguishes doublets in ImageStream data. (A) Cleanet doublet detection using five fluorescent channels, compared to manual doublet detection using two morphology channels. (B) Same comparison of Cleanet and manual doublet detection, overlayed on a UMAP plot of the five fluorescent channels. (C) The distribution of two membrane dyes and a viability dye in Raji cells, THP1 cells and the three doublet types they form. (D) Proportions of doublet types found by Cleanet (Observed) compared to expected proportions. In the sample treated with Rituximab, Cleanet finds a higher proportion of heterotypic doublets. (E) Brightfield images of five events sampled uniformly at random from those classified by Cleanet as singlets. (F) Brightfield images of five events sampled uniformly at random from those classified by Cleanet as doublets.

In the untreated sample, the observed doublet proportions matched the expected ones based on singlet proportions. But in the Rituximab treated sample, heterotypic doublet proportion was 0.15 higher than expected, and the homotypic proportions were correspondingly lower (Fig. 4D) This result serves as a proof of concept that Cleanet can be used to detect cell-cell interactions that are biological rather than technical in origin.

Finally, among all cells classified by Cleanet, five singlets and five doublets were sampled uniformly at random and the brightfield images for these events were inspected (Fig. 4E, 4F). All five images of putative singlets indeed showed a single cell. Meanwhile, four of the five images of putative doublets showed two or more cells, while the fifth was an edge case containing one clear cell surrounded by other fragments or debris. This result provides solid evidence that Cleanet makes accurate predictions and can be used to filter singlets with high specificity.

## Methods

### Data acquisition

Data was acquired as previously described for the mass cytometry dataset^23^, the flow cytometry dataset^18^, and the imaging cytometry dataset^22^. Manual gating annotations for the mass cytometry dataset are available with the data as deposited for the referenced work.

### Data preprocessing

Data was minimally processed before running Cleanet. For mass cytometry, bead removal and Gaussian gating was performed manually using the OMIQ platform (Supplementary Fig. 1A). This step removed relatively few events, so it was not strictly necessary. A helper function included in the Cleanet package was used to remove events whose distance from the origin in protein expression space was smaller than a threshold. For flow cytometry, a helper function included in the Cleanet package was used to remove events whose sum of FSC-A and SSC-A measurements is less than 50,000 units (Supplementary Fig. 2A). Both helper functions provide visual feedback and allow users to adjust the threshold to better fit debris in their data. Flow data was compensated, but no data transformation was applied to either mass or flow cytometry data before running Cleanet. ImageStream data was gated to remove debris and dead cells (Supplementary Fig. 3A).

### Cleanet: doublet detection

The doublet detection module of Cleanet is inspired by the DoubletFinder algorithm, previously published for use with scRNA-seq data. The steps of the doublet detection algorithm are:

1. Sample uniformly at random N/2 pairs of events, where N is the total number of events.
2. For each sampled pair, add the (untransformed) values of the corresponding two rows in the matrix of protein expression data. Call the resulting N/2 extra rows “simulated doublets”.
3. Create an augmented dataset containing the N original events and N/2 simulated doublets. Perform a hyperbolic arcsin transform on the augmented dataset, with cofactor provided as a parameter. (The default is 5 for mass cytometry and 500 for flow cytometry.)
4. Find the 15 nearest neighbors of each event in the augmented dataset. The Hierarchical Navigable Small Worlds method^24^ is used to find fast approximate nearest neighbors.
5. For each of the original N events, if more than 5 of its 15 nearest neighbors are simulated doublets, predict it is a doublet. Otherwise, predict it is a singlet.
6. Compute the confidence metric as the percentage of simulated doublets for which more than 5 out of 15 neighbors are simulated doublets.

The user can choose the markers to be considered by Cleanet, as well as the cofactor to be used for data transformation before computing nearest neighbors. The algorithm is applied independently to each file, so inter-sample variability does not affect the results.

### Cleanet: deconvoluting doublet composition

Cleanet can classify doublet composition if given as input the results of the doublet detection algorithm, as well as an array specifying the cell type of all singlet events. The algorithm is:

1. Determine the composition of each of the N/2 simulated doublets. For each simulated doublet, use the pair of indices from which it was constructed and return the cell type of the two components. For an index corresponding to a singlet, read off the cell type from the input array. For an index corresponding to a doublet, use the most frequent cell type among its 15 nearest neighbors computed during doublet detection.
2. To classify the composition of predicted doublets, use majority vote among the nearest neighbors which are simulated doublets, and thus already have a known composition from step 1. By construction, each predicted doublet has at least 5 simulated doublet neighbors.

### Expected frequency of doublet types

The expected frequency used in Fig. 3C and Fig. 4D is computed assuming that each pair of singlets is equally likely as any other pair to pass through the detector at the same time and be registered as a doublet. Given an array of cell type labels for singlets, the frequency of cell types is tabulated. Then the expected frequency of a doublet type (such as Monocyte-T cell) is the product of the frequencies of the component cell types (Monocyte and T cell), up to a normalization constant which is determined from the percentage of doublets in the sample. For heterotypic doublets, the order of components is irrelevant, so the frequencies for the two possible orders (Monocyte-T cell and T cell-Monocyte) are summed. Biological interaction between specific cell types could be one source of disagreement between expected and observed frequencies.

### Comparison with other methods

The RemoveDoublets function in the PeacoQC R package was used with its default settings: nmad=4, channels FSC-A and FSC-H.

For mass cytometry data, hierarchical manual gating was performed using the OMIQ platform, first to detect singlets and then to determine cell types. To make Figure 2C, cell type labels were transferred from the singlets determined by manual gating to the singlets determined by Cleanet, on a file by file basis. Since gate coordinates could not be exported from OMIQ, each gate was approximated in R as the convex hull of all events inside the gate. This set of approximate gates was then applied in hierarchical fashion to the singlets determined by Cleanet.

## Discussion

In this article, we introduced Cleanet, an automated method for doublet detection in cytometry data. To the best of our knowledge, this is the first time that a method based on multivariate protein expression has been used to detect doublets in cytometry. Thus, the main goal of this work is to demonstrate that multivariate cytometry data contains valuable information about doublets that is not necessarily visible in bivariate plots.

In our experiments, Cleanet showed greater than 90% agreement with manual gating or the existing method PeacoQC for most files under consideration. When Cleanet differed significantly from manual gating, the differences mostly reflected better performance for Cleanet in the context of poor DNA intercalator signal. Therefore, the main advantages of Cleanet are automation and robustness to the quality of individual channels. When using multiple channels, even if they are individually noisy, multivariate analysis can improve resolution, assuming that the noise in different channels is not too highly correlated.

Aside from detecting doublets, Cleanet can classify them by the cell type of their component singlets. As of this writing, it is uncommon to keep track of doublet types in cytometry data or explore their biological relevance. However, our method enables the identification of possible applications, such as measuring enrichment of T cell-monocyte complexes or other cell-cell interactions^16^. This manuscript provides a proof of concept, detecting a treatment-dependent increase in Raji-THP1 heterotypic doublet proportions.

Cleanet is robust to the choice of markers included in the analysis, as long as there are enough markers with good staining index. Intuitively, performance is better if there is an available lineage marker for each cell type present in the sample. Conversely, if some cells are negative for all markers, then doublets in which they participate may fail to be detected. Cleanet reports a confidence score with each run, which can be used to determine if enough markers have been included.

A caveat of the Cleanet approach is that debris events can create misleading simulated doublets: for example, a debris-T cell simulated doublet may be similar to a T cell singlet, and enough of these may lead the algorithm to classify T cell singlets as doublets. With small amounts of debris, this phenomenon is not enough for T cell singlets to pass the 5 out of 15 nearest neighbors criterion and be classified as doublets. But if debris comprises more than 20% of the sample, it should be depleted in a pre-processing step. The Cleanet package provides helper functions to deplete debris in either mass or flow cytometry before proceeding to doublet identification.

Another inconvenience is the running time of the nearest neighbor computation, which can be a few minutes for a sample of one million events, compared to less than a second for a bivariate gate. As a matter of efficiency, the best we can do is include the list of nearest neighbors with the other Cleanet output, for potential use by other downstream algorithms (for example, Phenograph).

As the first multivariate method to detect doublets in cytometry, we hope that Cleanet will help automate data analysis workflows, leading to more scalable and robust analysis results for immunology studies.

## Competing Interests

No competing interest is declared.

## Author Contributions Statement

## Acknowledgments

We thank Takuya Ohtani and the CyTOF Core at the University of Pennsylvania for mass cytometry data acquisition. This work benefited from ImageStreamX MarkII funded by NIH S10OD019942-01.

## Notes

### Competing Interest Statement

The authors have declared no competing interest.

