## Supplementary Material for "Cleanet: robust doublet detection in cytometry data based on protein expression patterns"

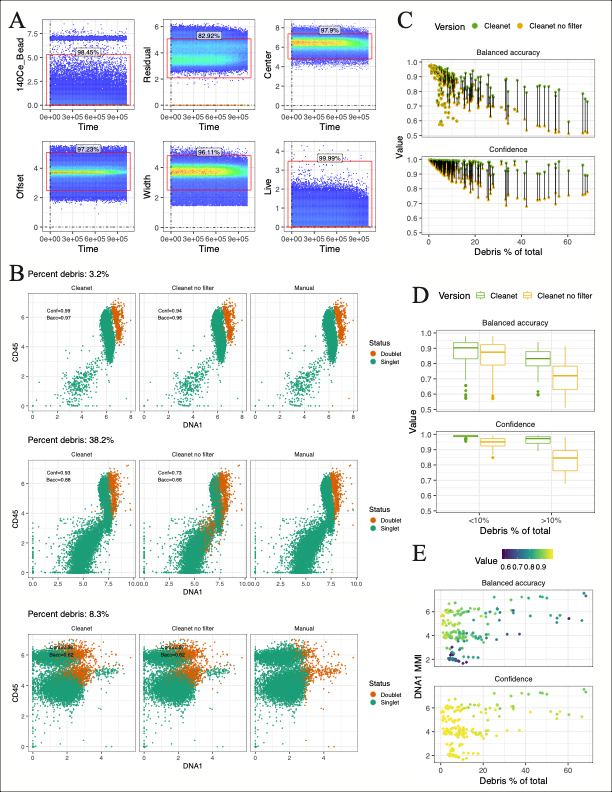

Supplementary Figure 1 Additional information and analysis for mass cytometry data. (A) Representative bivariate gating plots for the manual Bead, Gaussian and Viability gates that were applied to all mass cytometry files. (B) For each of three files, Cleanet predictions with or without debris filtering, compared with manual gating of doublets. Top row: a file with low debris proportion and good DNA intercalator staining. Middle row: a file with high debris proportion, impairing Cleanet simulation of doublets in the version with no filtering. Bottom row: a file with poor DNA intercalator staining, impairing accurate manual gating of doublets. (C) Accuracy metrics decrease with debris proportion, but debris filtering recovers much of the lost performance. (D) Impact of debris filtering on Cleanet performance is most notable for high debris proportion. (E) Cleanet diverges from manual gating for files with low median metal intensity (MMI) for DNA intercalator, indicating poor intercalator staining.

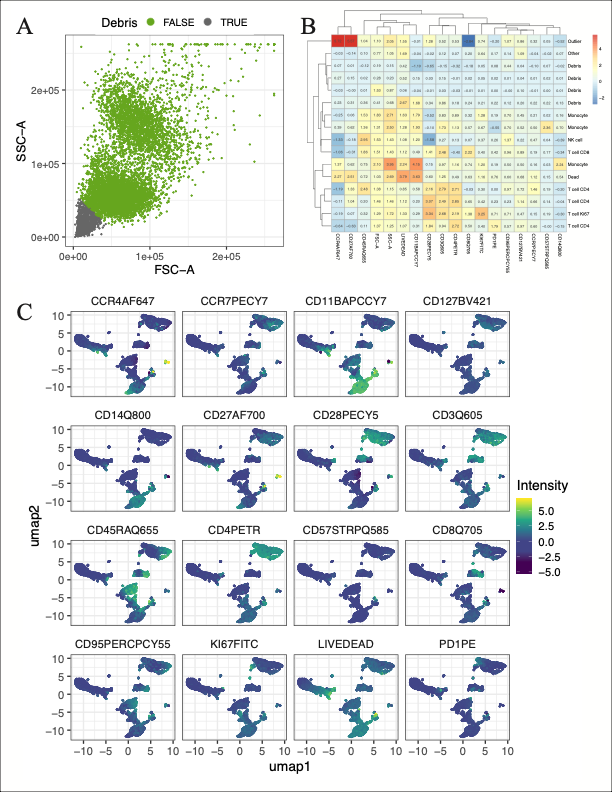

Supplementary Figure 2 Additional information and analysis for flow cytometry data (A) Debris depletion performed by Cleanet helper function, using default settings. (B) Heatmap showing manual annotation of Phenograph clusters based on mean fluorescence intensity. (C) UMAP dimensional reduction overlaid with fluorescence intensity for all 16 channels considered by Cleanet.

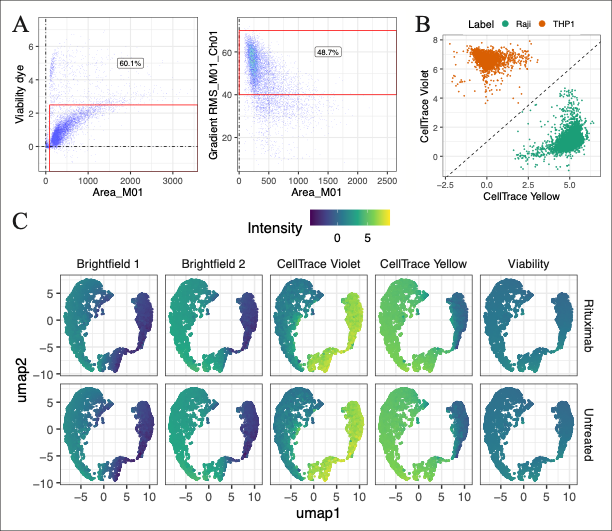

Supplementary Figure 3 Additional information and analysis for ImageStream data. (A) Successive gates used to select viable cells for analysis. (B) After Cleanet removed doublets, singlets were classified based on the two membrane dyes which labeled each cell type. (C) UMAP dimensional reduction overlaid with brightfield or fluorescence intensity for all 5 channels considered by Cleanet.
